# The Nicotinamide Salvage Pathway is a Metabolic Vulnerability of High-Risk MDS Stem Cells

**DOI:** 10.64898/2025.12.19.695528

**Authors:** Sweta B. Patel, Daniel Moskop, Steven Moriera, Stephanie Gipson, Colin C. Anderson, Alexendra Crook, Maxwell McCabe, Daniel Stephenson, Hannah E. Terry, Andrew Kent, Tracy Young, Anna Krug, Caitlin Price, Monica Ransom, Regan Miller, Ana Vujovic, Mohammad Minhajuddin, Mark J. Althoff, The University of Colorado Interdisciplinary Joint Biology Program (The CUIJBP Consortium), Anthony J. Saviola, Brett M. Stevens, Robert S. Welner, Ekaterina L. Andrianova, Andrei V. Gudkov, Travis Nemkov, Angelo D’Alessandro, Austin E. Gillen, Daniel A. Pollyea, Craig T. Jordan, Eric M. Pietras

## Abstract

High-risk myelodysplastic syndrome (HR-MDS) is a malignant clonal disorder originating in hematopoietic stem and progenitor cells (HSPCs). The current standard of care for HR-MDS patients is hypomethylating agents; however, the response rate is poor. There is thus a need to explore vulnerabilities of HR-MDS HSPCs for better clinical outcomes. We demonstrate that HR-MDS HSPCs have significant upregulation of metabolic proteins required for glycolysis, citric acid cycle, and oxidative phosphorylation. Consistently, we see increased oxygen consumption rate in HR-MDS HSPCs compared to healthy, suggesting an increased metabolic rate. Corroboratively, compared to healthy HSPCs, HR-MDS HSPCs have increased abundance of mitochondrial complex I proteins, which are NADH dehydrogenases, and crucial for energy production. Therefore, we investigated whether HR-MDS HSPCs are functionally reliant on NAMPT, the rate-limiting enzyme in the nicotinamide salvage pathway of NAD anabolism. NAMPT inhibition significantly decreased NAD(H) in HR-MDS HSPCs. Consequently, NAMPT inhibition reduced the oxygen-consuming capacity of HR-MDS-HSPCs compared to healthy. Importantly, NAMPT inhibition significantly impaired the self-renewal and colony-forming potential, increased cell death and reduced disease burden specifically of HR-MDS HSPCs, compared to healthy controls. Collectively, our data suggest that NAMPT is selectively required for the function and survival of HR-MDS HSPCs representing a promising therapeutic target.

## INTRODUCTION

High-risk myelodysplastic syndrome (HR-MDS) is a malignant blood disorder characterized by increased blasts, severe cytopenias, and dysplasia of hematopoietic lineages in the bone marrow (BM). HR-MDS often presents with recurrent cytogenetic and mutational lesions and carries an increased risk of progression to acute myeloid leukemia (AML)(1, 2). Monotherapy with hypomethylating agents like azacitidine and decitabine is the current standard of care for HR-MDS. However, only 17% of the HR-MDS patients achieve complete remission, and the median overall survival from diagnosis is 18.6 months(3, 4). Furthermore, the majority of the HR-MDS patients are elderly and often unfit for hematopoietic stem cell transplant, which remains the only curative treatment(5). Thus, new therapeutic options are needed to improve outcomes of HR-MDS patients.

Biologically, HR-MDS originates in the hematopoietic stem and progenitor cell (HSPC) population residing in the BM and is marked by CD34 surface expression(6–9). Importantly, these cells also constitute a persistent reservoir contributing to disease relapse and, in many cases, progression to AML. We and others have previously shown that inhibition of protein synthesis can selectively induce cell death in HR-MDS HSPCs(9–13). Furthermore, we found that inhibition of protein synthesis reduces cellular respiration in HR-MDS blasts(9), suggesting oxidative phosphorylation could be a critical downstream target of protein synthesis inhibition in HR-MDS HSPCs. This indicates a link between the metabolic properties of HR-MDS HSPCs and AML stem cells, which also exhibit reliance on oxidative phosphorylation for cellular energy production and survival(14, 15).

Rate-limiting enzymes required to facilitate oxidative phosphorylation-dependent metabolism in malignant leukemia stem cells(16–18) use nicotinamide adenine dinucleotide (NAD) as a co-factor. As NAD is consumed, it is broken down to nicotinamide, which enters the nicotinamide salvage pathway to be recycled back into NAD(19). Compared to other pathways that make NAD, like the de novo and Preiss-Handler pathways, the salvage pathway is more energy efficient at maintaining stable NAD pools, and hence is selectively utilized by malignant cancer cells(19–21). The rate-limiting enzyme for the nicotinamide salvage pathway is NAMPT (nicotinamide phosphoribosyltransferase), which has been demonstrated as a potential therapeutic target in relapse/refractory AML and patients with TP53 deficiency and/or loss of chromosome 7(22–26), as well as in other malignancies(27).

Here, we address the importance of NAMPT and NAD in the function and survival of HR-MDS HSPCs. We employed mass spectrometry-based metabolomics and proteomics to determine differences between HR-MDS and healthy HSPCs. We demonstrated that HR-MDS HSPCs have higher oxygen consumption compared to normal and an increased abundance of metabolic proteins that use NAD as a co-factor. Subsequently, we inhibited NAMPT using pharmacological and genetic approaches to reduce NAD, which consequently impaired oxidative phosphorylation in HR-MDS HSPCs by reducing carbon flux through glycolysis and oxidative citric acid cycle. Furthermore, we utilized stem and progenitor cell functional assays, like colony forming and engraftment potential, to decipher the impact of genetic and pharmacological NAMPT inhibition on HR-MDS HSPCs. Lastly, we employed *in vivo* xenografts to explore the significance of NAD metabolism on HR-MDS disease burden. Altogether, we find that NAMPT inhibition not only impaired the function of HR-MDS HSPCs compared to normal but also induced cell death and reduced disease burden. These data identify NAMPT inhibition as a therapeutic target for HR-MDS HSPCs.

## RESULTS

### HR-MDS HSPCs have enriched metabolic proteins compared to healthy control HSPCs

We have previously shown that inhibiting protein synthesis using omacetaxine preferentially eliminates HR-MDS HSPCs compared to healthy control HSPCs(9, 13). Therefore, to identify differences in the proteome of HR-MDS and healthy control HSPCs (CD45+ Lineage-CD34+ surface expression), we conducted LC-MS-based proteomics analysis. HR-MDS HSPCs were isolated from patients with IPSS-M scores ranging from -0.28 to 4.27 and IPSS-R scores from 4.9 to 8, which are considered high to very-high risk MDS (Fig 1A, Table S1)(2). Further, to encompass generalizable aspects of HR-MDS biology, we employed specimens bearing mutations in splicing factors (SF3B1, SRSF2, U2AF1, ZRSR2), epigenetic modifiers (TET2, DNMT3a, ASXL1, EZH2, IDH2, STAG2), transcription factors (GATA2, ETV6, WT1, SETBP1, CEBPA, TP53) and signaling genes (NF1, CBL, CSF3R, NRAS, KRAS) (Fig 1A, Table S1). 5 of the 12 patients used in this study also carried cytogenetic abnormalities common to MDS such as (del)5q, -7, (del)7q or +8, while 4 patients had a complex karyotype (Fig 1A, Table S1).

**Figure 1:**
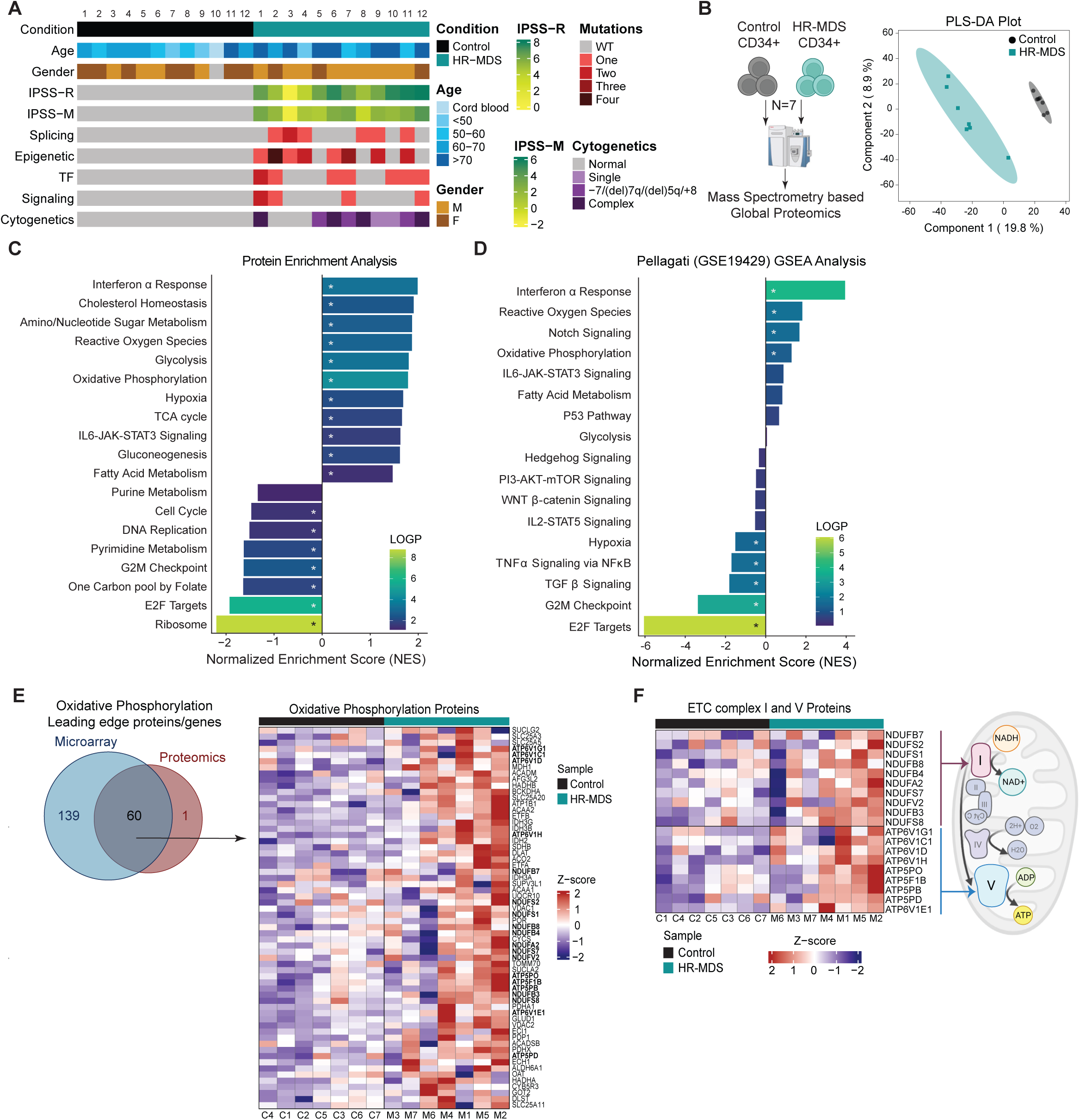
HR-MDS HSPCs have enriched metabolic proteins compared to healthy control HSPCs. **(A)** Clinical characteristics and associated metadata of patient and healthy control samples used in the study, including age, gender, IPSS scores, cytogenetics and mutational profile. For cytogenetics, patients are classified into 3 groups, those having a) normal karyotype; b) common score driving abnormalities, like loss of chr7, (del)7q, (del)5q, and/or gain of chr8; c) other single cytogenetic abnormalities; and/or d) complex karyotype. For mutational profile, patients are grouped based on presence of one-four a) Splicing mutations, which include SF3B1, SRSF2, U2AF1, ZRSR2; b) epigenetic modifier mutations which include TET2, DNMT3a, ASXL1, EZH2, IDH2, STAG2; c) transcription factor mutations which include GATA2, ETV6, WT1, SETBP1, CEPBA, TP53; and/or d) signaling mutations which include NF1, CBL, CSF3R, NRAS, KRAS. Specific details for each patient are provided in Supplemental Table 1. **(B-C)** PCA plot (B) and Protein enrichment analysis (C) of LC/MS-based global proteomics comparing HR-MDS (N=7) to healthy control (N=7) CD34+ HSPCs. Specimens used – M1-M7 and C1-C7. **(D)** GSEA analysis of GSE19429 microarray data comparing HR-MDS (N=183) to healthy control (N=17) CD34+ HSPCs. Star represents significant pathways. **(E)** Venn diagram depicting leading-edge proteins and genes from Oxidative phosphorylation (OxPhos) pathway of the proteomic and microarray GSEA, respectively. Heatmap comparing HR-MDS (N=7) with healthy control (N=7) CD34+ HSPCs for the abundance of the 60 common Oxidative phosphorylation (OxPhos) proteins. Specimens used – M1-M7 and C1-C7. **(F)** Heatmap comparing abundance of the electron transport chain complex I and complex V proteins between HR-MDS (N=7) and healthy control (N=7) CD34+ HSPCs. Specimens used – M1-M7 and C1-C7.

Our proteomics analysis was carried out on HSPCs isolated from 7 HR-MDS patients versus 7 age- and sex-matched healthy control BM specimens, which identified 540 differentially expressed proteins between HR-MDS and healthy control HSPCs, with 228 proteins upregulated (fold change≥1.2; p≤0.05) and 312 proteins downregulated (fold change≤0.8; p≤0.05) in HR-MDS HSPCs relative to healthy controls. Partial Least Squares-Discriminant Analysis (PLS-DA) analysis identified a 20% variance across Principal Component 1 (PC1), which comprised of changes in the abundance of ribosomal and metabolic proteins between the HR-MDS and healthy control HSPC proteomes (Fig 1B). Protein enrichment analysis of differentially expressed proteins revealed that HR-MDS HSPCs had an enrichment for metabolic pathways like nucleotide metabolism, hypoxia, reactive oxygen species, glycolysis, oxidative phosphorylation, and citric acid cycle (Fig 1C). Additionally, HR-MDS HSPCs were negatively enriched for mRNA translation and ribosomal pathways (Fig 1C).

To validate our findings, we analyzed publicly available Affymetrix GeneChip RNA microarray data (GSE19429) comparing HSPCs from 183 HR-MDS patients with 17 healthy controls(28). Consistent with our data, gene set enrichment analysis of published transcriptomic data showed that compared to healthy control HSPCs, HR-MDS HSPCs likewise had significantly enriched metabolic pathways, specifically reactive oxygen species, oxidative phosphorylation, and fatty acid metabolism (Fig 1D). Notably, the leading-edge oxidative phosphorylation proteins identified from our proteomics data almost completely overlapped with leading-edge genes from the published microarray data (Fig 1E). Additionally, 20 of the 60 overlapping genes and proteins constituted subunits of mitochondrial complex V (ATP synthase, the final step of oxidative phosphorylation that converts ADP to ATP) and complex I (NADH dehydrogenase, which oxidizes NADH generated by glycolysis and citric acid cycle to NAD+) of the electron transport chain (ETC) (Fig 1F). Collectively, these data indicate that HR-MDS HSPCs are primed at gene expression and protein levels to support oxidative metabolism.

### HR-MDS HSPCs have increased oxidative phosphorylation

Given the increased abundance of metabolic genes and proteins in HR-MDS HSPCs compared to healthy control HSPCs, we sought to identify whether these changes in expression are accompanied by increased metabolic activity in HR-MDS HSPCs. Consequently, we sorted and expanded HSPCs from 3 HR-MDS and 3 healthy BM specimens to measure oxygen consumption rates using Seahorse, which provides kinetic measurement of oxidative metabolism. Indeed, HR-MDS HSPCs demonstrated elevated maximal respiration and spare respiratory capacity in HR-MDS compared to healthy control HSPCs (Fig 2A).

**Figure 2:**
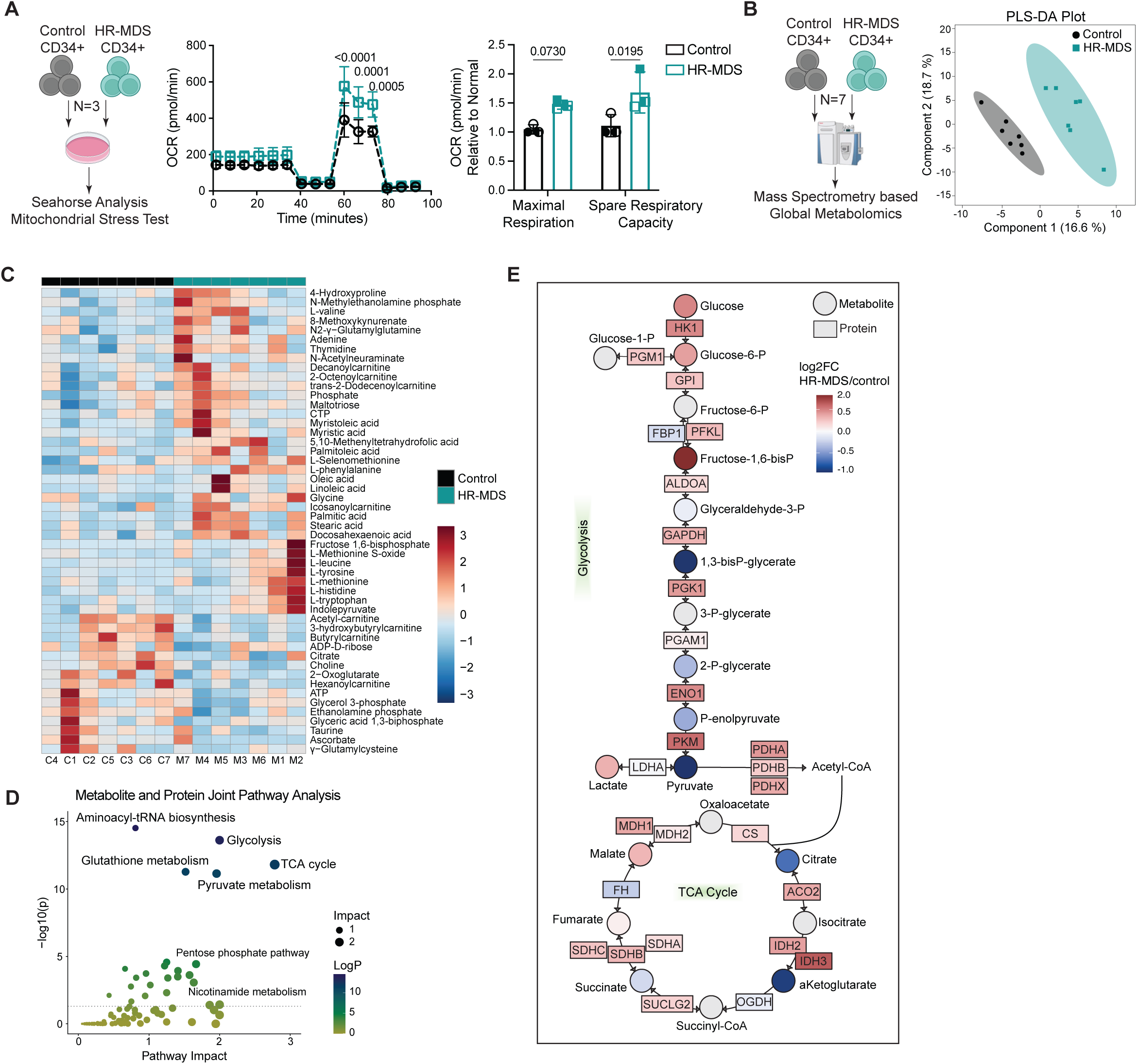
HR-MDS HSPCs have increased oxidative phosphorylation. **(A)** Mitochondrial Stress Test seahorse analysis on expanded CD34+ HSPCs from HR-MDS (N=3) and healthy control (N=3) BM specimens. Specimens used – M5, M7-M8 and C7, C11-12. **(B)** PCA plot of UHPLC/MS-based global metabolic profiles comparing HR-MDS (N=7) and healthy control (N=7) CD34+ HSPCs. Specimens used – M1-M7 and C1-C7. **(C)** Heatmap comparing top 50 differentially abundant metabolites between HR-MDS (N=7) and healthy control (N=7) CD34+ HSPCs. Specimens used – M1-M7 and C1-C7. **(D)** Scatter plot depicting results of joint pathway analysis of the metabolomic and proteomic data using Metaboanalyst, comparing differentially expressed proteins and metabolites in HR-MDS versus healthy control CD34+ HSPCs. Pathway impact scores is normalized degree centrality (number of links connecting a node) topology measures of genes/metabolites in each pathway(51). **(E)** Summary of metabolic and protein changes in central carbon metabolism in HR-MDS versus healthy control HSPCs. Proteins are indicated by rectangles, and metabolites are indicated by circles. The filled color of each is proportional to the log2(fold change) of HR-MDS CD34+ HSPCs with respect to healthy control CD34+ HSPCs, where blue to red represents a log2(fold change) of −1 to 2, respectively. Gene symbols are provided for each protein. ACO2, aconitase; ALDOA, aldolase; CS, citrate synthase; ENO1, enolase; FBP1, fructose-1,6-bisphosphatase; FH, fumarate hydratase; GAPDH, glyceraldehyde-3-phosphate dehydrogenase; GPI, glucose-6-phosphate isomerase; HK1, hexokinase; IDH, isocitrate dehydrogenase; LDHA, lactate dehydrogenase-A; MDS, malate dehydrogenase; OGDH, oxoglutarate dehydrogenase; PDH, pyruvate dehydrogenase; PFKL, phosphofructokinase, liver type; PGAM1, phosphoglycerate mutase; PGK1, phosphoglycerate kinase; PGM1, phosphoglucomutase; PKM, pyruvate kinase; SDH, succinate dehydrogenase; SUCLG2, succinyl-CoA synthetase.

To further characterize metabolic differences between HR-MDS HSPCs and healthy control HSPCs, we performed unbiased global metabolomic analysis(29) on HSPCs isolated from 7 HR-MDS and 7 healthy BM specimens using ultra high-performance liquid chromatography coupled to mass spectrometry (UHPLC-MS). PLS-DA analysis showed a 16% variance across PC1 between the groups (Fig 2B), which was comprised of changes in abundance of amino acids, central carbon and one-carbon metabolites, and fatty acids. These differences were also evident in the analysis of the top 50 differentially expressed metabolites (Fig. 2C, Fig S2A), which revealed increased abundance of amino acids and fatty acids, and reduced abundance of portions of the citric acid cycle, short chain acylcarnitines, and lipid headgroup intermediates in HR-MDS HSPCs (Fig 2C).

To understand the interplay between changes in metabolic protein expression and the corresponding metabolite level in HR-MDS HSPCs, we next performed joint pathway analysis of both datasets. We identified that citric acid cycle and glycolysis were the most significant alterations in HR-MDS HSPCs (Fig 2D), characterized by elevated expression of metabolic proteins with concomitant modulation in the steady state abundance of metabolites along these pathways (Fig 2E). Taken together, these data support a model in which elevated expression of metabolic proteins facilitates increased flux through glycolytic and central carbon pathways, resulting in elevated oxidative metabolism.

### NAMPT is essential to maintain increased metabolic activity in HR-MDS HSPCs

Oxidative metabolism is coupled to the ETC, which consumes oxygen and requires electrons provided by reducing equivalents NADH and FADH to produce energy. HR-MDS HSPCs have increased abundance of ETC complex I (Fig 1F), which oxidizes NADH to NAD+ to provide electron for energy production. In addition, our proteomic analysis identified an increased abundance of proteins and metabolic enzymes that maintain NAD/NADH redox balance, use NAD as their cofactor, or are reliant on NAD for their function. These proteins include glucose-6-phosphate dehydrogenase (G6PD), malate dehydrogenase (MDH1), malic enzyme (ME2), isocitrate dehydrogenase (IDH2), NAD kinase 2 (NADK2), and sirtuin 2 (SIRT2) (Fig 3A)(30). This result suggested that HR-MDS HSPCs might require an increased supply of cellular NAD to support elevated oxidative metabolism. Interestingly, our UHPLC-MS analysis did not identify any differences in the overall abundance of NAD+, NADH, or the ratio of oxidized/reduced (NAD+/NADH) NAD in HR-MDS versus healthy control HSPCs (Fig 3B-C). We thus reasoned that HR-MDS HSPCs would be uniquely reliant on the salvage pathway to rapidly recycle NAD in support of increased NAD reliant enzymes and an elevated oxidative metabolism.

**Figure 3:**
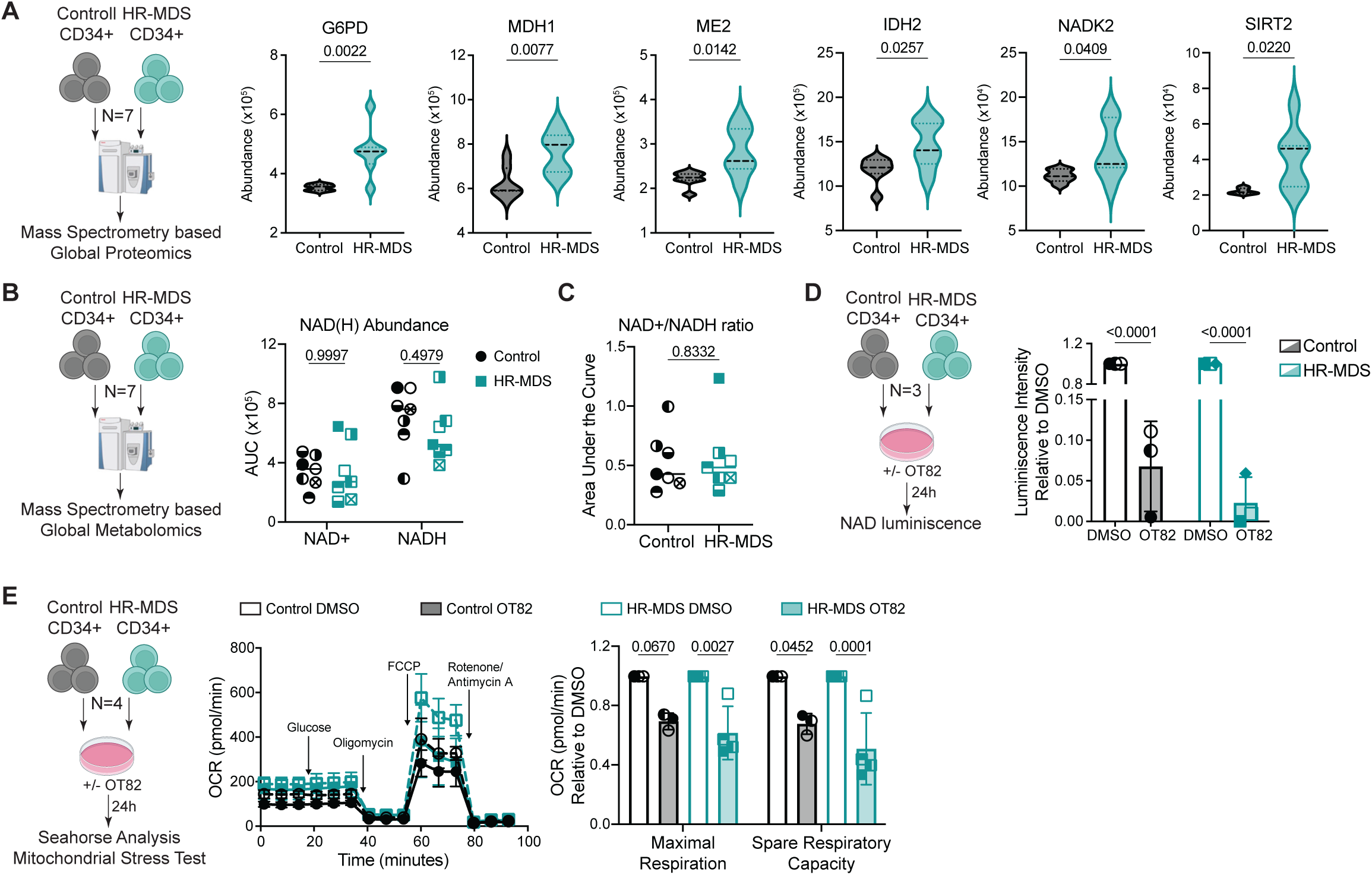
NAMPT is essential to maintain increased metabolic activity in HR-MDS HSPCs. **(A)** Violin plots comparing abundance of G6PD, MDH1, ME2, IDH2, NADK2, and SIRT2 between HR-MDS (N=7) and healthy control (N=7) CD34+ HSPCs measured using LC/MS. Specimens used – M1-M7 and C1-C7. **(B-C)** Abundance of NAD+ and NADH (B) and ratio of NAD+/NADH (C) measured using UHPLC/MS in HR-MDS (N=7) and healthy control (N=7) CD34+ HSPCs. Specimens used – M1-M7 and C1-C7. **(D)** NAD level measured using luminescence in HR-MDS (N=3) and healthy control (N=3) CD34+ HSPCs after 24h of treatment with DMSO or 100nM OT-82. Specimens used – M3, M7, M9 and C5-C7. **(E)** Mitochondrial Stress Test seahorse analysis on HR-MDS (N=4) and healthy control (N=3) CD34+ HSPCs after 24h of treatment with DMSO or 100nM OT-82. Specimens used – M5-M8 and C7, C11-12.

To address this hypothesis, we chose to pharmacologically inhibit NAMPT, the rate-limiting enzyme of the salvage pathway, using OT-82(31) due to its higher potency and specificity compared to other NAMPT inhibitors(20, 32). We first verified the effect of OT-82 on NAD levels using MDS-L cells, a cell line generated from the BM of a HR-MDS patient(33), and found that OT-82 completely abrogated cellular NAD pools by 24h using an orthogonal NAD luminescence assay (Fig S3A). We further evaluated the impact of NAMPT inhibition with OT-82 on cellular respiration in MDS-L cells using the Seahorse analyzer. We observed a significant reduction in both maximal and spare respiratory capacity, indicating that NAMPT inhibition leads to reduced oxidative metabolism (Fig S3B). Given these results, we next assessed the impact of NAMPT inhibition on the metabolic activity of primary HR-MDS HSPCs. Like MDS-L cells, we found that OT-82 efficiently depleted NAD pools in HR-MDS HSPCs as well as in healthy control HSPCs (Fig. 3D). However, NAMPT inhibition triggered a more significant reduction in maximal respiration and spare respiratory capacity of HR-MDS HSPCs (Fig. 3E). Altogether, these data indicate that HR-MDS HSPCs are preferentially reliant on the nicotinamide salvage pathway and NAD for their metabolic activity compared to healthy control HSPCs.

### NAMPT inhibition disrupts central carbon metabolism in HR-MDS HSPCs

To further understand how NAMPT inhibition impacts metabolism in HR-MDS HSPCs, we carried out both steady state metabolomics and stable isotope-labeled metabolite tracing assays, with or without NAMPT inhibition using MDS-L cells as a surrogate for HR-MDS HSPC. We treated MDS-L cells ± 100nM OT-82 for 24h and first performed global metabolic profiling using UHPLC-MS. PLS-DA analysis identified a 47% variance in metabolites across PC1 between OT-82 and DMSO-treated MDS-L (Fig 4A). As expected, NAMPT inhibition significantly reduced nicotinamide levels in addition to other nucleotides, like ADP, AMP (Fig 4B, S4A). Likewise, we observed a significant depletion of ATP, specifically in OT-82-treated HR-MDS HSPCs compared to healthy control HSPCs (Fig S4B). Furthermore, OT-82 reduced the levels of citric acid cycle metabolites, fatty acids, and acyl-carnitines while causing accumulation of amino acids and glycolysis intermediates in MDS-L cells (Fig S4A). Likewise, NAMPT inhibition triggered significant increases in the abundance of upper glycolytic intermediates glucose-6-phosphate and fructose 1-6-bisphosphate, with corresponding reductions in the lower glycolysis metabolite 2/3-phosphoglycerate, along with a relative increase in the non-oxidative pentose phosphate pathway (PPP) intermediate sedoheptulose 7-phosphate and decrease in citrate, in HR-MDS HSPCs versus healthy control HSPCs (Fig. S4C). Collectively, these data suggest that NAD depletion caused by NAMPT inhibition impaired mitochondrial metabolism leading to a rerouting of carbon towards glycolysis and tangential pathways such as the PPP.

**Figure 4:**
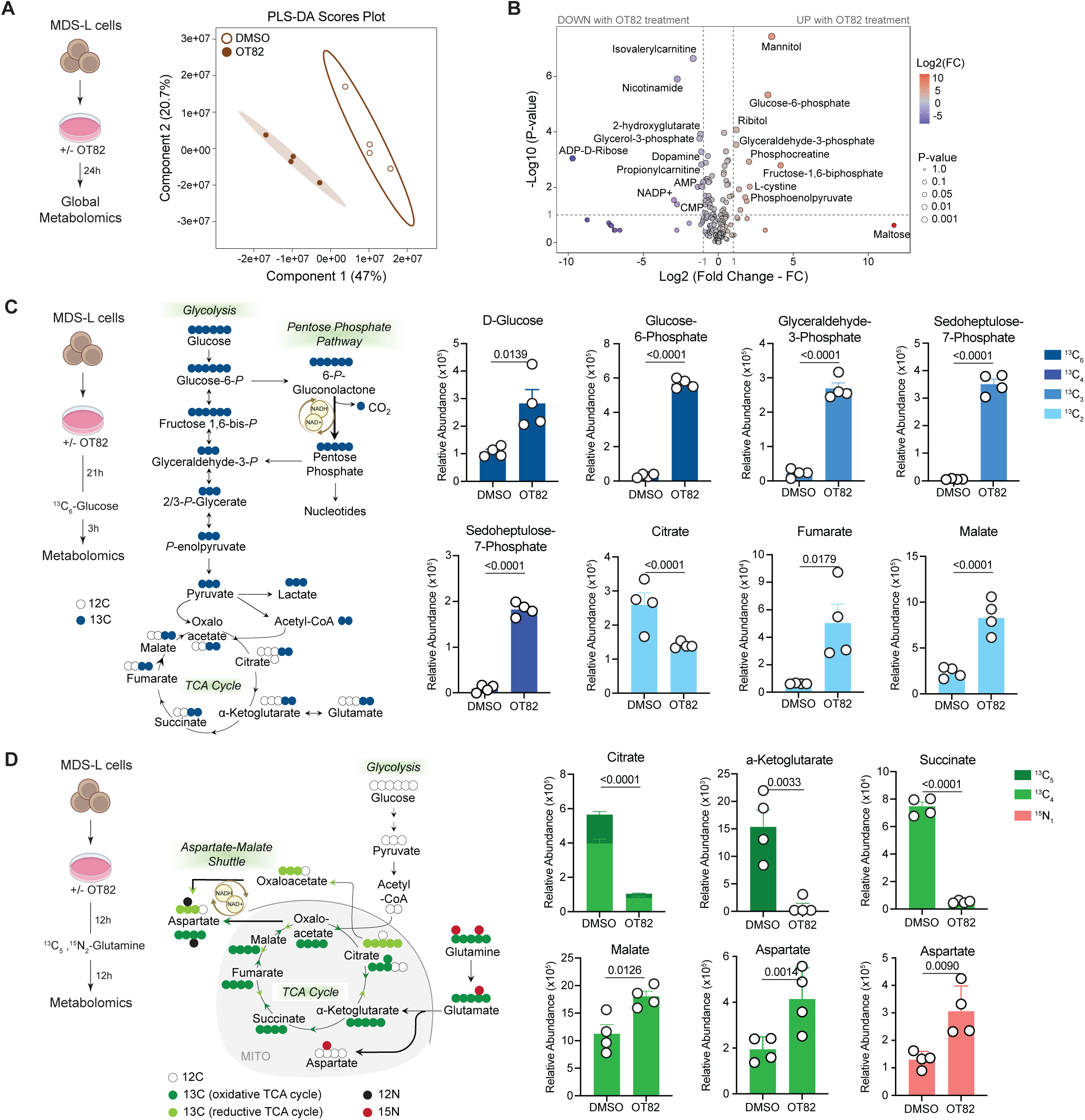
NAMPT inhibition disrupts central carbon metabolism in HR-MDS HSPCs. **(A-B)** PCA plot (A) and Volcano plot (B) of UHPLC/MS-based global metabolic profiles comparing MDS-L cells treated with DMSO or 100nM OT-82 for 24h. **(C)** Abundance of ^13^C labeled metabolites in glycolysis and TCA cycle post 3hr of ^13^C_6_-glucose incubation in MDS-L cells treated with DMSO or 100nM OT-82 for 24h total. The metabolites were measured with UHPLC/MS. **(D)** Abundance of ^13^C labeled metabolites in oxidative and reductive TCA cycle and aspartate-malate shuttle post 12hr of ^13^C_5_^15^N_2_-glutamine incubation in MDS-L cells treated with DMSO or 100nM OT-82 for 24h total. The metabolites were measured with UHPLC/MS.

To validate these findings, we treated MDS-L cells with 100nM OT-82 for 24h and in the last 3h of treatment added ^13^C_6_-glucose to trace the fate of glucose in the context of NAMPT inhibition (Fig 4C). Consistent with our global metabolomics analysis (Fig S4A), OT-82 treatment significantly increased abundance of ^13^C labeled isotopologues of glucose, glucose-6-phosphate and glyceraldehyde-3-phosphate (Fig 4C). Interestingly, this was accompanied by a dramatic increase in ^13^C labeling of the PPP intermediate sedoheptulose-7-phosphate (Fig. 4C), suggesting increased PPP cycling to boost NADPH regeneration and compensate for NAD depletion. Furthermore, we observed a significant reduction in ^13^C-labeled citrate (Fig 4C), indicative of reduced pyruvate entry into the TCA cycle via pyruvate dehydrogenase. However, increased ^13^C_3_ labeled malate (Fig 4C), suggests enhanced activity of malic enzyme in OT-82-treated cells. In addition, a more pronounced increase of ^13^C_2_ malate and fumarate, in line with decreased citrate labeling, suggest a re-routing of carbon that favors aspartate aminotransferase activity and the malate-aspartate shuttle to replace lost intermediates and boost NAD(H) reconversion(34).

To assess whether NAMPT inhibition activates the malate-aspartate shuttle pathway, we performed a similar isotope labeling experiment using ^13^C5,^15^N2-L-Glutamine in MDS-L cultures in the last 12h of OT-82 treatment (Fig 4D). Mass spectrometry analysis revealed that NAMPT inhibition reduced the flux of carbons from labeled glutamine in citrate, succinate, and α-Ketoglutarate (Fig 4D). On the other hand, we observed increased flux of glutamine-derived carbons and nitrogen into malate (^13^C_4_ label) and aspartate (^15^N_1_ and ^13^C_4_ labels) (Fig 4D). These results suggest that the HR-MDS HSPCs activate specific metabolic pathways like PPP and the malate-aspartate shuttle to compensate for reduced NAD levels resulting from NAMPT inhibition (Fig 3D, S3A). This implies that NAD is an important substrate for HR-MDS HSPCs, forcing cells to activate compensatory pathways to make up for the loss of NAMPT-derived NAD.

### Nicotinamide metabolism is essential for the survival and function of HR-MDS HSPCs

To determine the functional role of NAMPT in survival, we treated HSPCs from HR-MDS and healthy control specimens with 100nM OT-82 for 24-48h and analyzed cell death using Annexin V flow cytometry analysis. After 48h of NAMPT inhibition, HR-MDS HSPCs had a significant 3-fold increase in apoptosis and a 50% reduction in total viable cell count, while the healthy control HSPCs were not significantly impacted (Fig 5A, S5A-C). This impact on apoptosis was completely rescued by supplementing the MDS-L cells with NAD (Fig S5D), implying reliance of HR-MDS HSPCs on NAD. We further sought to validate NAMPT as a functional target using a CRISPR-based knockout (KO) approach in MDS-L cells. We confirmed complete KO of *NAMPT* by demonstrating the lack of expression of NAMPT protein via Western blot (Fig S5E). KO of NAMPT not only reduced NAD levels as measured by luminescence (Fig S5F) but also reduced the survival capability of the MDS-L as compared to the AAVS1-KO control MDS-L, as read out by selective loss of the GFP+ *NAMPT*-deficient cells versus GFP+ AAVS1-targeted cells (Fig 5B, S5G).

**Figure 5:**
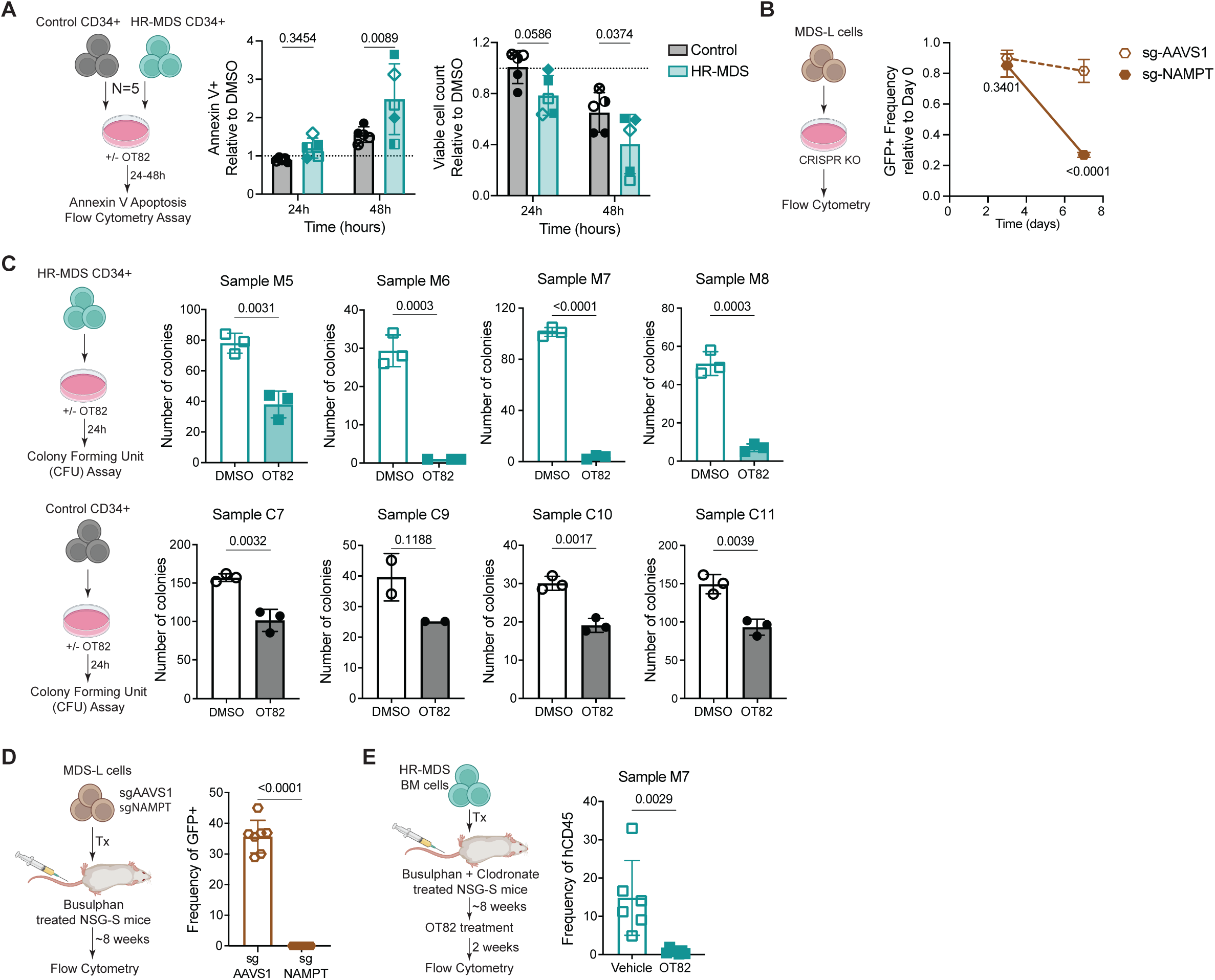
Nicotinamide metabolism is essential for the survival and function of HR-MDS HSPCs. **(A)** Frequency of Annexin V+ cells normalized to DMSO (left) and number of viable cells normalized to DMSO (right) measured using flow cytometry for HR-MDS (N=5) and healthy control (N=5) CD34+ HSPCs post 24-48h of treatment with DMSO or 100nM OT-82. Each point is a biological replicate, and the dotted line represents normalized DMSO value of 1. Specimens used – M6-M7, M9-12 and C3, C6-C9. **(B)** Frequency of GFP+ AAVS1 and NAMPT knockout MDS-L cells measured over time at day 3 and day 7 using flow cytometry. **(C)** Number of colonies counted at day 12 for HR-MDS (N=4) and healthy control (N=4) CD34+ HSPCs plated into CFU media post 24h of treatment with DMSO or 100nM OT-82. Specimens used – M5-M8 and C7, C9-C11. **(D)** Frequency of GFP+ hCD45 MDS-L cells in BM of NSG-S mice xenografts transplanted with AAVS1 and NAMPT knockout MDS-L cells. **(E)** Frequency of hCD45+ cells in BM of NSG-S mice xenografted with primary HR-MDS patient BM sample and treated with 25mg/kg OT-82 OG for 2 weeks (5 days on, 2 days off). Specimen used – M7.

We next assessed the impact of NAMPT inhibition on the function of HR-MDS HSPCs using colony-forming unit (CFU) assay to measure clonogenic activity. Except for one HR-MDS specimen, which showed a 50% reduction in colony-forming potential, NAMPT inhibition using 100nM OT-82, almost completely abrogated the colony-forming potential of HR-MDS HSPCs (Fig 5C, S5H). This contrasted with normal HSPCs, which retained 70% colony-forming capability (Fig 5C, S5H) at the same concentration of OT82. This suggests that NAMPT inhibition preferentially targets the function of HR-MDS HSPCs. Furthermore, to determine the importance of NAMPT on the engraftment potential of HR-MDS HSPCs, an essential parameter of stem cell function, we transplanted AAVS1 and NAMPT-KO MDS-L cells into NSG-S mice. Strikingly, *NAMPT*-KO MDS-L cells completely lost their engraftment potential as observed by the absence of GFP+ *NAMPT*-KO hCD45+ cells in the NSG-S mice as compared to 40% engraftment of GFP+ AAVS1-KO hCD45+ cells, with no change in total marrow count (Fig 5D, S5I-J).

Additionally, to address whether NAMPT inhibition targets HR-MDS HSPCs *in vivo*, we transplanted MDS-L cells as well as HR-MDS patient BM into NSG-S mice. Post engraftment, the mice were treated with 25mg/kg OT-82 for 2 weeks(32). Noticeably, flow cytometry analysis upon treatment completion revealed a significant reduction in xenografted MDS-L cells (Fig S5K) and nearly complete eradication of the primary HR-MDS graft (Fig 5E, S5M) with no change in total marrow count (Fig S5L and S5N). Conversely, OT-82 treatment of healthy C57/B6 mice using the same treatment regimen had no impact on the abundance of BM HSPCs, while only a minimal change in the distribution of BM myeloid and lymphoid cells (Fig S5O-P). Moreover, OT-82 treatment did not impact the long-term multilineage repopulating capacity of the BM of these mice when transplanted into busulfan-conditioned recipients and analyzed 16 weeks later (Fig. S5Q). Altogether, these data indicate that HR-MDS HSPCs have a preferential reliance on NAMPT for their function and survival as opposed to normal HSPCs, thus providing a therapeutic vulnerability for HR-MDS patients.

## DISCUSSION

HR-MDS is a heterogenous malignant clonal disorder of the hematopoietic stem cells. Since the approval of hypomethylating agents in 2004, which are the current standard of care, no new treatment modalities have been approved for HR-MDS(5, 35). Hence, HR-MDS still remains an incurable disorder with only 1.7 years of median overall survival(2). The only curative option is allogeneic hematopoietic stem cell transplant, however, many HR-MDS patients are ineligible due to comorbidities, leaving non-transplant treatment approaches as the only option. There is thus an unmet clinical need to identify better therapeutic targets to improve HR-MDS patient outcome.

To this end, we and others have previously shown that HR-MDS is vulnerable to inhibition of protein synthesis(9–13). In the present study, we sought to evaluate the unique proteome of HR-MDS HSPCs, the source of the disease, to understand HR-MDS HSPC biology with the goal of identifying specific targetable vulnerabilities. We demonstrate that HR-MDS HSPCs have an increased abundance of metabolic proteins compared to healthy control HSPCs. Indeed, we show that HR-MDS HSPCs have an enhanced metabolic rate and oxygen consumption compared to their normal counterparts. This is consistent with previous characterizations of AML stem cells, which have identified the reliance of AML stem cells on oxidative phosphorylation as a targetable metabolic vulnerability, giving rise to the use of the BCL2 inhibitor venetoclax, IDH1 inhibitor ivosidenib and IDH2 inhibitor enasidenib in AML patients unfit for bone marrow transplant(14, 15, 36–39). Importantly, the metabolic features of HR-MDS HSPCs remain incompletely characterized, with the relatively broad spectrum of disease pathogenesis and mutational/karyotype heterogeneity of the HSPC compartment likely contributing to the current paucity of analysis. Further studies are warranted to draw mechanistic links between the MDS-HSPC proteome and cellular metabolism.

Accordingly, our data identified increased rates of oxidative phosphorylation and increased expression of metabolic proteins requiring NAD as a key feature that distinguishes HR-MDS HSPC from healthy controls. Despite this, we observed no corresponding change in the size of the NAD/H pool, which led us to suspect that HR-MDS HSPCs could be increasingly reliant on rapidly recycling NAD via the nicotinamide salvage pathway to sustain activity of the proteins that use NAD for their function, as well as the elevated metabolic activity observed in these cells. Indeed, we find that inhibition of NAMPT, negatively impacts the oxygen consumption rate, specifically of HR-MDS HSPCs. Our team previously showed that NAMPT inhibition in AML favored metabolic rewiring via activation of fatty acid desaturases, to compensate decreases in NAD pool with increased NADH recycling(23). Here, we show that HR-MDS cells activate the malate-aspartate shuttle and PPP as compensatory mechanisms, upon NAMPT inhibition. The malate-aspartate shuttle can facilitate TCA cycle anapleurosis and cytosolic NADH reoxidation. This is consistent with studies of T cells that have limited oxidative phosphorylation activity due to impaired mitochondrial fusion, which also activate the malate-aspartate shuttle and PPP at the expense of maximal carbon oxidation, underscoring the interplay between mitochondrial architecture, NAD homeostasis, and salvage dependence(40).

Additionally, our data show that despite increased activation of these compensatory pathways, neither can sustain cellular energetics and functional activity of HR-MDS HSPCs upon the loss of NAD caused by NAMPT inhibition, triggering their selective killing, similar to AML stem cells(22–26). A major challenge in the treatment of HR-MDS is the capacity of the malignant stem cells to evolve and trigger disease relapse and/or progression to AML. NAMPT itself may represent a crucial metabolic pressure point for malignant cells at multiple stages of disease pathogenesis, as it plays a critical role in recycling NAD for use in oxidative metabolism. As recent work has defined increased reliance on oxidative phosphorylation to be a feature of pre-leukemic cells harboring mutations in TET2 and DNMT3A, reliance on NAMPT likewise may be established early in pathogenesis(41, 42). These data together suggest that the requirement for NAMPT may represent a stage-specific feature of myeloid pathogenesis. Future studies can establish whether reliance on NAMPT is associated with disease progression from pre-leukemia to MDS to AML.

In summary, our data indicate that NAMPT, the rate-limiting enzyme for the nicotinamide salvage pathway of NAD anabolism, is important for maintenance of oxidative phosphorylation and is critical for the survival of HR-MDS HSPCs. Inhibition of NAMPT uniquely eliminates HR-MDS HSPCs versus healthy control HSPCs, indicating this enzyme represents a unique therapeutic vulnerability that can be exploited for improved treatment of HR-MDS. Our pre-clinical work thus provides strong support for the initiation of NAMPT inhibitor clinical trials to improve outcomes for high-risk MDS patients.

## MATERIALS and METHODS

### Sex as a biological variable

Our study examined male and female patients as well as male and female animals, and similar findings are reported for both sexes.

### Human Specimens

Primary human HR-MDS specimens were obtained from BM aspirates at diagnosis and normal healthy specimens were obtained from femoral head of people undergoing hip replacement surgery at the University of Colorado. All patients gave written informed consent for procurement of samples on the Institutional Review Board (IRB) protocol of the University of Colorado (IRB 12-0173 and IRB 20-1908) and the University of Alabama at Birmingham (IRB X160420008) in accordance with an assurance filed with and approved by the Department of Health and Human Services. All specimens were acquired in accordance with recognized ethical guidelines of the Declaration of Helsinki and U.S. Common Rule. Details on age, gender, cytogenetics, mutation, IPSS-R, and IPSS-M score. are included in Suppl Table 1.

### Cell Culture

#### Human Specimen Culturing

Primary high-risk MDS and healthy control CD34+ cells were cultured using SFEM II media (Stem Cell Technologies, 09655) supplemented with human cytokines – 25ng/mL SCF (PeproTech, 300-07), 25ng/mL TPO (PeproTech, 300-18), 10ng/mL IL3 (PeproTech, 200-03), 50ng/mL FLT3 (PeproTech, 300-19) and 0.5uM SR1 (Stem cell technology, 72342), and 1uM UM729 (Stem cell technology, 72332)(43, 44). For expansion of CD34+ cells, media was changed every 3 days and cells stepped up in well size upon reaching confluency.

#### Cell line culturing

MDS-L cell line was a generous gift from Daniel Starczynowski and cultured with RPMI media supplemented with 10% FBS, 10ng/mL human IL3 and 1% antibiotic-antimycotic.

#### Drug treatment

OT-82 was manufactured and provided by Drs. Ekaterina L. Andrianova and Andrei V. Gudkov at Roswell Park(32). *In vitro* cells were treated with drug at 100nM dose for 24-72h as stated in the figure legends.

### Cell Sorting

Primary human high-risk MDS and normal specimens were sorted for CD34+ stem and progenitor cells. Briefly, specimens were thawed and stained with Live/dead viability dye (EMD Millipore, 278298; dilution 500 nmol/L) to exclude dead cells, CD45 (BD, 563716; dilution 1:100), Lineage cocktail (BioLegend, 348701; dilution 1:100), CD34 (BD, 562577; dilution 1:100) and CD38 (BD, 560677; dilution 1:100) to identify stem and progenitor cells(6–8).

### Microarray Data Analysis

Microarray gene expression data from high-risk MDS patients and healthy controls were obtained from GEO dataset GSE19429, comprising Affymetrix Human Genome U133 Plus 2.0 arrays. The raw expression data were log2 transformed, probe identifiers were mapped to gene symbols using the hgu133plus2.db annotation package (v3.13.0), and inter-array normalization was performed using quantile normalization. Differential gene expression analysis was performed using the limma package(45) (v3.60.6) with linear models fitted to MDS patients versus healthy controls.

Gene set enrichment analysis was performed with CAMERA(46) (v1.60.0) using Hallmark and Gene Ontology Molecular Function gene sets from MSigDB (v25.1.1). All analyses were performed in R (v4.4.0).

### Proteomics

#### Sample preparation

120,000 primary CD34+ cells from healthy control and HR-MDS BM specimens were washed with ice-cold PBS and pellets flash frozen. Samples were submitted for metabolite/protein extraction and liquid chromatography coupled to mass spectroscopy (LC-MS)-based analysis.

#### Extraction of proteins

Digestion was performed using ProtiFi S-Trap™ micro spin columns (Prod #: C02-micro-80) according to the manufacturer’s protocol. The samples were suspended in 50 μL PBS and 50 μL 2x lysis buffer (10% SDS in 100 mM triethylammonium bicarbonate (TEAB)). The samples were then vortexed for 10 minutes and centrifuged at 21,000 x g for 5 minutes. The desired supernatant was pulled into a separate tube, and the remaining samples were stored. The 100 μL solution was used for a standard S-trap micro spin column protocol with accommodating reagents. Samples were reduced with 4.3 ul 100 mM tris (2-carboxyethyl) phosphine (TCEP) and incubated at 55 °C for 15 minutes. The samples were then alkylated by adding 4.3 ul 500 mM-chloroacetamide (CIAA) and incubated at room temperature in the dark for 10 minutes. The reduced and alkylated samples then received 10.9 ul 27.5% phosphoric acid to a final concentration of 10% and 717 μL wash/binding buffer. The solution was transferred to the S-Trap micro column and centrifuged 4,000 x g for 30 seconds. The columns were washed 6 times with 150 μL wash/binding buffer. Five of the six washes were centrifuged at 4,000 x g for 30 seconds, and the final wash was spun for 90 seconds. The columns were then transferred, and 20 μL 0.05 μg/μL trypsin buffer was added for an overnight digest at 37C. The samples were eluted the next day via 50 mM TEAB, 0.2% FA, and 50% acetonitrile (ACN); 40 ul elution for each. The eluted digests were dried down in a speed vacuum and rehydrated in 75 μL 0.1% formic acid (FA). Samples were cleaned for mass spectrometry analysis utilizing Pierce C18 Spin Columns (Prod #: 84850). Briefly, samples were loaded onto spin columns and washed with 0.1% FA to remove contaminants. Samples were eluted off the column with 80% acetonitrile in 0.1% FA, dried down, and resuspended in 0.1% FA at a concentration of 0.025 μg/μL.

#### High-throughput Mass Spectrometry

Data generated using a timsTOF mass spectrometer (Bruker Daltonics) coupled with an Evosep One liquid chromatography-mass spectrometry (LC-MS) interface. In brief, digested peptides were loaded onto Evotips following the manufacturer’s protocol and analyzed directly using an Evosep One liquid chromatography system (Evosep Biosystems, Denmark). Peptides were separated on a 75 µm i.d. × 15 cm separation column packed with ReproSil 1.9 µm C18 beads, 120A resin (Evosep Biosystems, Denmark), and over a predetermined 44-minute gradient according to manufacturer settings. LC mobile phase solvents consisted of 0.1% formic acid in water (buffer A) and 0.1% formic acid in 80% acetonitrile (buffer B, Optima LC/MS, Fisher Scientific). The system was coupled to the timsTOF Pro mass spectrometer (Bruker Daltonics, Bremen, Germany) via the nano-electrospray ion source (Captive Spray, Bruker Daltonics). Instrument control and data acquisition were performed using Compass Hystar (version 6.0) with the timsTOF Pro (Bruker Daltonics, Bremen, Germany) operating in parallel accumulation-serial fragmentation (PASEF) mode. The following parameters were utilized for data-dependent acquisition in positive ion mode: mass range 100-1700m/z, ion mobility scan from 0.7 to 1.40 Vs/cm^2^, ramp accumulation times were 100ms; capillary voltage was 1400 V, dry gas 3.0 L/min and dry temp 180°C. The PASEF settings were as follows: 10 MS/MS scans (total cycle time, 1.1 s); charge range 0 – 5; active exclusion for 0.40min; scheduling target intensity 12500 cts/s; intensity threshold 1000 cts/s; collision-induced dissociation (CID) energy 20 eV to 59 eV.

#### Data analysis

Raw data were searched using MSFragger(47) (version 4.1) through the FragPipe platform (version 22.0). Precursor tolerance was set to ± 15 ppm, and fragment tolerance was set to ± 0.08 Da, allowing for 2 missed cleavages. Data was searched against the Human SwissProt sequence database (UP000005640) with common contaminants and decoys added. Trypsin semi-specific cleavage was utilized, with fixed modifications set as carbamidomethyl (C) and variable modifications set as oxidation (M) and acetyl (protein N-term). Results were filtered to 1% false discovery rate (FDR) at the peptide and protein level using Philosopher employed through FragPipe. Label-free quantification (LFQ) was performed using the IonQuant (version 1.10.27), including unique and razor peptides. Protein intensities were sum normalized, log transformed and auto-scaled in MetaboAnalyst 6.0 (https://www.metaboanalyst.ca/)(48). One factor Statistical analysis was used for volcano plots and PLS-DA figure generation. Enrichment analysis was carried out in R, using Kegg and Hallmark datasets.

### Metabolomics

#### Sample preparation

For global metabolomics, 100,000 MDS-L cells and 20,000 primary CD34+ cells from healthy control and HR-MDS bone marrow specimens were treated with DMSO and 100nM OT-82 for 24h. For carbon flux studies, 500,000 MDS-L cells were treated with DMSO and 100nM OT-82 for 24h. At the final 3h and 12h, 2.5mM heavy labeled glucose (^13^C_6_; Cambridge Isotope Laboratories, CLM-1396-1) and 0.5mM heavy labeled glutamine (^13^C_6_,^15^N_2_; Cambridge Isotope Laboratories, CNLM-1275-H-0.1) were spiked into the culture respectively.

Next day, cells were washed with ice-cold PBS and counted and pellets flash frozen in quadruplicates (triplicates for primary specimens). Cells were submitted for metabolite extraction and ultra high-performance LC-MS (UHPLC-MS)-based analysis.

#### Extraction of metabolites

Cells were extracted to a final concentration of 2E06 per mL of cold MeOH:ACN:H2O (5:3:2, v:v:v). For low cell count samples (primary specimens), 150 uL of buffer was added. Samples were then vortexed at 4 °C for 30 minutes. Following vortexing, samples were centrifuged at 12700 RPM for 10 minutes at 4 °C and supernatant was transferred to a new autosampler vial for analysis. Low cell-count samples were dried down and reconstituted in 35 µL of buffer before injection. A portion of extract from each sample was also combined to create a technical mixture, injected throughout the run for quality control.

#### High-throughput Metabolomics Analysis

Analyses were performed as previously published via a modified gradient optimized for the high-throughput analysis of metabolomics(29). Briefly, the analytical platform employs a Vanquish UHPLC system (Thermo Fisher Scientific, San Jose, CA, USA) coupled online to an Exploris 120 mass spectrometer (Thermo Fisher Scientific, San Jose, CA, USA). Metabolomics extracts were resolved over an ACQUITY UPLC BEH C18 column (2.1 x 100 mm, 1.7 µm particle size) held at 45 °C (Waters, MA, USA). For positive mode mobile phase (A) 0.1% FA in water and mobile phase (B) 0.1% FA in ACN was used. For negative mode mobile phase (A) 10mM Ammonium Acetate in Water and mobile phase (B) 10mM Ammonium Acetate in 50:50 ACN:MeOH was used. For negative and positive mode analysis, the chromatographic gradient was as follows: 0.45 mL/min flowrate for the entire run, 0% B at 0 min, 0% B at 0.5 min, 100% B at 1.1 min, 100% B at 2.75min, 0% B at 3min, 0% B at 5 min. For positive ion mode, full MS was utilized from 65 to 975 m/z at 120,000 resolution, with 3.5 kV spray voltage, 50 sheath gas, 10 auxiliary gas. For negative ion mode, full MS mode was utilized from 65 to 975 m/z at 120,000 resolution, with 3.4 kV spray voltage, 50 sheath gas, 10 auxiliary gas. Calibration was performed prior to analysis using the PierceTM Positive and Negative Ion Calibration Solutions (Thermo Fisher Scientific).

#### Data analysis

Raw data files were converted to .mzXML using RawConverter. Metabolites were assigned and peaks were integrated using Maven (Princeton University) in conjunction with the KEGG database and an in-house standard library. Peaks were integrated using Maven (Princeton University). Raw peak data for global metabolomics of MDS-L was normalized by mean-scaling and that for primary specimens was normalized to normal and auto-scaled in MetaboAnalyst 6.0 (https://www.metaboanalyst.ca/)(48). One factor Statistical analysis was used for volcano plots and PLS-DA.

### PDX engraftment

NSG-S mice were conditioned with 100uL clodronate liposomes (5mg/mL concentration obtained from liposoma) and 25 mg/kg busulfan (Thermo Scientific Chemicals, J61348) via i.p. injection 48h and 24h prior to transplant respectively(9, 49). The busulfan stock was made fresh at 25 mg/ml in 100% DMSO, the stock was then diluted 1:10 in pre-warmed saline (0.9% NaCl) down to 2.5 mg/ml right before use. The diluted busulfan solution was kept in 37°C water bath before IP injection to prevent precipitation of busulfan due to low solubility. Primary HR-MDS bone marrow cells were sorted on viability and resuspended in PBS supplemented with 2% FBS. Anti-human CD3 antibody (BioXCell) was added at a final concentration of 1μg/10^6^ cells to prevent potential graft-versus-host disease. Each mouse recipient received 0.1mL containing 1x10^5^ cells via tail vein injection; there were 8 to 10 mice per experiment group. Post 15-20% engraftment, mice were treated with 25 mg/kg OT-82 for 2 weeks (5 days ON, 2 days OFF) via oral gavage(32). Mice were sacrificed after 4 to 8 weeks and analyzed by flow cytometry for human CD45, CD34, CD38 and annexin V. All animal studies were done at the University of Colorado under Institutional Animal Care and Use Committee-approved protocol no. 308. The University of Colorado is accredited by the Association for Assessment and Accreditation of Laboratory Animal Care, abides by the Public Health Service Animal Assurance of Compliance, and is licensed by the United States Department of Agriculture.

### Colony-forming Assays

Human CD34+ cells from HR-MDS and healthy control BM specimens were isolated via FACS sorting, treated with DMSO and OT-82 for 24h, washed off drug, and added into methylcellulose (R&D systems, HSC003). 1000 CD34+ cells were plated per replicate in triplicates for each condition. Colonies were counted 10-12 days after initial plating.

### Seahorse Assay

500,000 expanded primary CD34+ cells obtained from healthy control and HR-MDS BM or MDS-L cells were treated with DMSO and 100nM OT-82 for 24 hours. Cells were harvested and resuspended in RPMI seahorse media with 2mM Glutamine and 1mM Pyruvate. 100K cells were plated per well in quadruplicate of a XFe96 plate coated with CellTak (Corning, 354240). Oxygen consumption rate (OCR) was measured at basal and after injection of 10mM Glucose, 5ug/ml oligomycin, 2uM FCCP and 5uM Antimycin A plus 5uM Rotenone on Seahorse XFe 96 Extracellular Flux Analyzer.

### Annexin V Flow

10,000 primary CD34+ cells from healthy control and HR-MDS bone marrow or MDS-L cells were treated with DMSO and 100nM OT-82 for 24-72h in triplicates. Cells were collected and stained with Annexin V (BD, 556421; dilution 1:100) and Live/dead viability dye (EMD Millipore, 278298; dilution 500 nmol/L) in 1X Annexin V buffer for 15min at room temperature. Cells were washed, resuspended in 1X Annexin V buffer to acquire on flow analyzer. To obtain cell counts, 10uL of counting beads (BioLegend, 424902) per replicate was used and calculation performed as per manufacturer instructions.

### NAD luminescence

10,000 primary CD34+ cells from healthy control and HR-MDS bone marrow and MDS-L cells were treated with DMSO and 100nM OT-82 for 24h in triplicates. Next day, cells were collected and resuspended in PBS. Subsequently, Promega NAD/NADH-Glo kit (G9071) was used as per manufacturer instructions.

### CRISPR knock-out (KO)

NAMPT KO MDS-L cells were prepared by CRISPR/Cas9 gene editing technology. DNA oligonucleotides harboring the forward and reverse sequences of the 20bp guide RNA, not including the PAM sequence, were ordered through IDT (https://idtdna.com). sgRNA was designed to target NAMPT (sg2: CTGGGAATGACAAAGCCCTC – targeting exon 4) using Broad CRISPick. sgRNA against AAVS1 (GGGGCCACTAGGGACAGGAT) safe harbor site was used as a control. The forward and reverse oligonucleotides were annealed to generate a double-stranded DNA fragment with 4bp overhangs compatible with ligation into BsmBI-digested LRG2.1 eGFP CRISPR-Cas9 plasmid (Addgene, #108098)(50). 293T Lenti-X (Takara Bio, 632180) were transfected using PEI with 60ug pLRG2.1 eGFP sgNAMPT.2 or sgAAVS1 along with 30ug psPAX2 (Addgene, #12260) lentiviral packaging plasmid and 15ug pMD2.G (Addgene, #12259) VSV-G envelope expressing plasmid to produce lentiviral particles. 1 million MDS-L cells were transduced separately with LRG2.1 eGFP sgAAVS1 and sgNAMPT.2 lentivirus overnight. After 24hrs fresh MDS-L media was added. KO was validated using western blot. eGFP was measured by flow cytometry 3-7 day’s post transduction for *in vitro* studies. For xenograft studies, 1 million KO cells were transplanted per NSG-S mouse.

### Immunoblotting

Cells were lysed using Pierce RIPA buffer (Thermo Scientific, 89900) supplemented with protease and phosphatase inhibitor (Thermo Scientific, 1861281) according to manufacturer’s instructions. Lysates were subsequently denatured using BME (Bio-Rad, 1610710) added to sample buffer laemmli (Bio-Rad, 1610747) followed by boiling for 10 minutes. Protein lysates were loaded into a polyacrylamide gel and transferred to a polyvinylidene difluoride (PVDF) membrane using the mini trans-blot transfer system (Bio-Rad). Membranes were probed with primary antibodies of the following targets: NAMPT (Cell Signaling Technologies, #61122; dilution 1:1000), B-Actin (Cell Signaling Technologies, #8457; dilution 1:1000). Membranes were incubated with primary antibodies overnight at 4°C on a shaker. After overnight incubation, membranes were incubated with respective horseradish peroxidase-conjugated secondary antibodies for 1 hour at room temperature. Subsequently, blots were exposed to SuperSignal West Femto Maximum Sensitivity Substrate (Thermo Scientific, 34096) and imaged using the ChemiDoc Imaging System (Bio-Rad). Images were processed using Image Lab.

### Statistical Analysis

All figures are mean ± SD of a representative data of at least 2-3 independent experiments or mean ± SD of at least 2-3 independent experiments. For each experiment, there were at least technical triplicates for every condition. Statistical analyses were performed depending on the spread of the variable. A Shapiro-Wilk test was used to determine normal versus abnormal distributions, and all continuous variables were tested for mean differences. Depending on the spread of variable both nonparametric: Mann–Whitney U test, ANOVA Kruskal-Wallis test, Wilcoxon test, and parametric: Unpaired student’s t-test and ANOVA were used. For ANOVA, Tukey’s or Sidak post-test was used to compare individual groups (GraphPad Prism version 10.0, La Jolla, CA).

### Data availability

All the raw proteomic and metabolomics data files are available via ProteomeXchange Consortium with identifier PXD070384 and the Metabolomics Workbench under DataTrack ID 6608. Any additional information required to reanalyze the data reported in this paper and requests for resources and reagents should be directed to and will be fulfilled by, Eric M. Pietras.

## AUTHOR CONTRIBUTIONS

Conceptualization, S.B.P., C.T.J., and E.M.P.; Data curation, S.G., C.C.A., A.C., M.MC., D.S., A.K., M.R., A.S., R.S.W., A.D., T.N., and A.E.G.; Analysis, S.B.P., S.G., C.C.A., A.C., M.MC., D.S., H.E.T., C.P., and M.R.; Funding acquisition, S.B.P., B.M.S., C.T.J., and E.M.P.; Investigation, D.M., S.M., C.C.A., A.C., M.MC., D.S., H.E.T., T.Y., A.K., C.P., R.M., A.V., M.M., and M.J.A.; Methodology, S.B.P., S.G., C.C.A., A.C., M.MC., D.S., T.N., A.D.; Project Administration, S.B.P.; Resources, S.G., A.K., T.Y., A.K., CUIJBP, A.S., R.S.W., E.L.A., A.V.G., A.D., B.M.S., D.A.P., C.T.J., and E.M.P; Supervision, A.S., A.D., T.M.N., A.E.G., C.T.J., and E.M.P.; Validation, S.B.P., A.S., A.D., T.M.N., A.E.G., C.T.J., and E.M.P.; Writing – original draft, S.B.P., S.M., C.C.A., A.C., D.S.; Writing – review and editing, S.B.P., A.V., M.J.A., A.J.S., B.M.S., R.S.W., A.V.G., T.N., A.D., D.A.P., C.T.J., E.M.P. All the authors read and approved the manuscript.

## ACKNOWLEDGEMENT

We would like to thank Drs. Cailin Collins, Mariya Amaya, James DeGregory, Zachary Walker, Hsin-Ying and the patients and their family members who contributed samples to our biobank, for their support. We also appreciate the contribution of femoral head specimens collected by The University of Colorado Interdisciplinary Joint Biology Program (The CUIJBP Consortium), consisting of Drs. Larry W. Moreland, Jennifer A. Seifert, Andrew D. Clauw, Prabil Kaini, Michael R. Dayton, Craig A. Hogan, Michael R. Clay, Michael J. Zuscik, Patrick M. Carry, V. Michael Holers, Daniel K. Moon, Cheryl L. Ackert-Bicknell and Anna Helena Jonsson.

