## Supplemental Data for "The Nicotinamide Salvage Pathway is a Metabolic Vulnerability of High-Risk MDS Stem Cells"

**Supplemental Table 1: HR-MDS patient and Healthy control metadata**

| MDS Patient ID | Age/Gender | IPSS-R | IPSS-M | Cytogenetics | Mutations |
|---|---|---|---|---|---|
| M1 | 70/M | 6.5 - very high | 3.14 - very high | deletion TP53 gene at 17p13.1,3 copies PML gene at 15q24 | CBL, CEBPA, KRAS, TET2, WT1 |
| M2 | 61/F | 5 - high | 1.89 - very high | normal | SF3B1, TET2, ASXL1, CSF3R, CEBPA, STAG2, EZH2 |
| M3 | 82/M | 0 - low | neg1.74 - moderate | normal | SRSF2, ZRSR2, TET2 |
| M4 | 36/M | 4 - intermediate | 0.95 - high | normal | SRSF2, ASXL1, IDH2 |
| M5 | 83/F | 6.26 - high | 0.65 - high | Gain of Chr 8 | None detected by qPCR |
| M6 | 76/M | 7.5 - very high | 2.66 - very high | loss of chr7 19, 22; del of 5q13; add 16q22, 3p3, 11p15 | TP53, TET2 |
| M7 | 58/M | 6.29 - very high | 3.07 - very high | Loss of Chr 7 | GATA2, TET2, EZH2, ETV6, NF1, ASXL1 |
| M8 | 66/M | 4.9 - high | neg0.28 - moderate | Gain of 11q23, 17p13, 20q11.2 | SF3B1 |
| M9 | 74/M | 6.08 - very high | 2.13 - very high | Gain of 1q21 sequences | IDH2, ASXL1, U2AF |
| M10 | 74/M | 7.55 - very high | 1.79 - very high | Rare cells with gain of 11q23 | U2AF1, WT1 |
| M11 | 57/M | 7.87 - very high | 2.55 - very high | Loss of Chr 7 | DNMT3A, SETBP1, SRSF2, TET2 |
| M12 | 70/F | 8 - very high | 4.27 - very high | Loss of 5q31, Gain of 6p22, Loss of 7q31 | NRAS, TP53 |

| Healthy Control ID | Age/Gender | History |
|---|---|---|
| C1 | 62/F | N/A |
| C2 | 51/F | N/A |
| C3 | 64/M | Cardiovascular Hx |
| C4 | 67/F | Cardiovascular Hx, former smoker |
| C5 | 68/M | Cardiovascular Hx, former smoker |
| C6 | 59/M | Cardiovascular Hx |
| C7 | 45/F | Former smoker |
| C8 | 56/F | N/A |
| C9 | 21/M | Former smoker |
| C10 | Cord blood | N/A |
| C11 | 72/F | Cardiovascular Hx, former smoker |
| C12 | 64/F | Cardiovascular Hx, Inflammatory polyarthropathy |

### SUPPLEMENTAL FIGURES

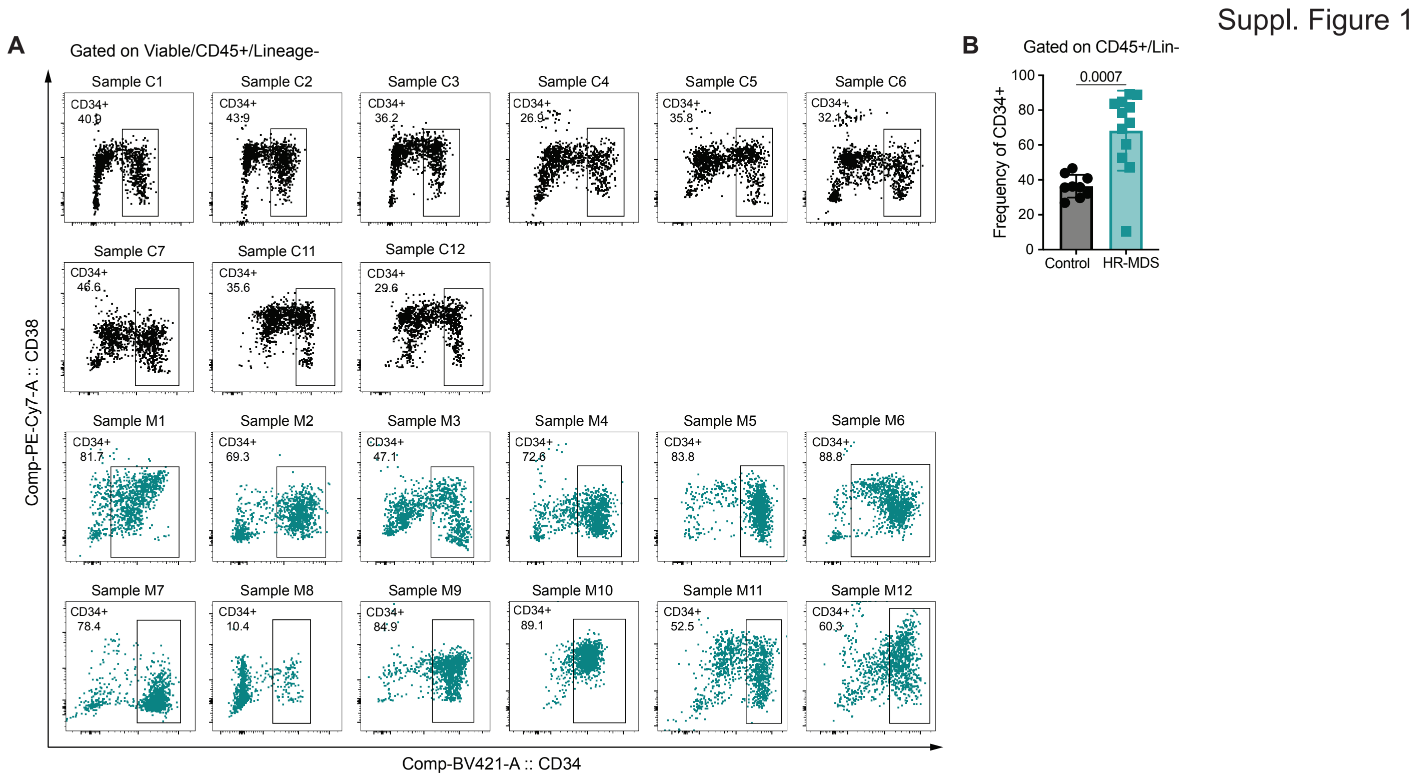


**Supplemental Figure 1: Immunophenotypic representation of HR-MDS and healthy control HSPCs used in the study**

**(A)** Flow plots showing CD34 and CD38 surface expression of representative healthy control (N=9) and all HR-MDS patient (N=12) samples used in the study. Cells are gated on viable, CD45+ and lineage-. Specimens showed – M1-M12 and C1-C7, C11-C12.

**(B)** Frequency of CD34+ cells in representative healthy control (N=9) and all HR-MDS patient (N=12) samples used in the study. Cells are gated on viable, CD45+ and lineage-. Specimens showed – M1-M12 and C1-C7, C11-C12.

A

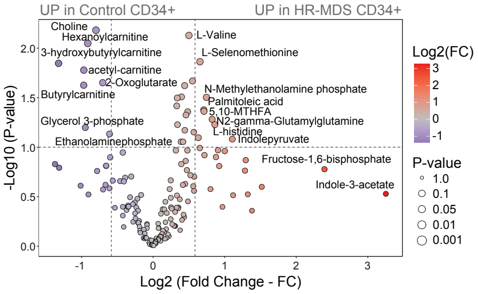


**Supplemental Figure 2: HR-MDS HSPCs have a distinct metabolic profile compared to healthy control HSPCs**

**(A)** Volcano plot of UHPLC/MS-based global metabolic profiles comparing HR-MDS (N=7) and healthy control (N=7) CD34+ HSPCs. Specimens used – M1-M7 and C1-C7.

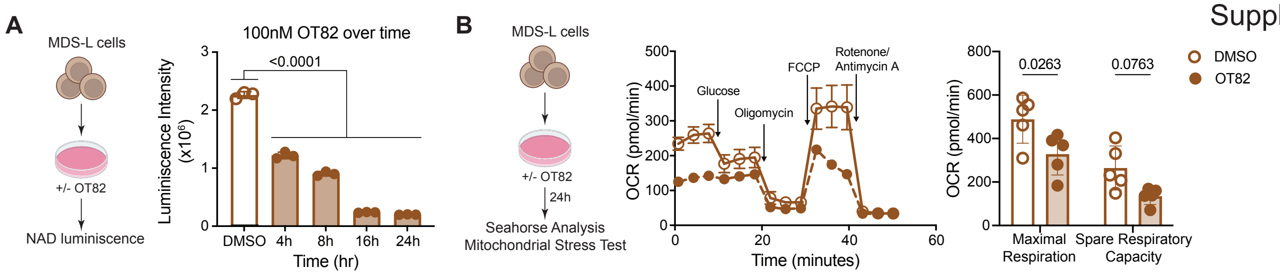


**Supplemental Figure 3: OT-82 reduces NAD and oxygen consumption in MDS-L cells**

**(A)** NAD level measured in MDS-L cells after 4-24h of treatment with DMSO or 100nM OT-82 using luminescence.

**(B)** Mitochondrial Stress Test seahorse analysis on MDS-L cells treated with DMSO or 100nM OT-82 for 24h.

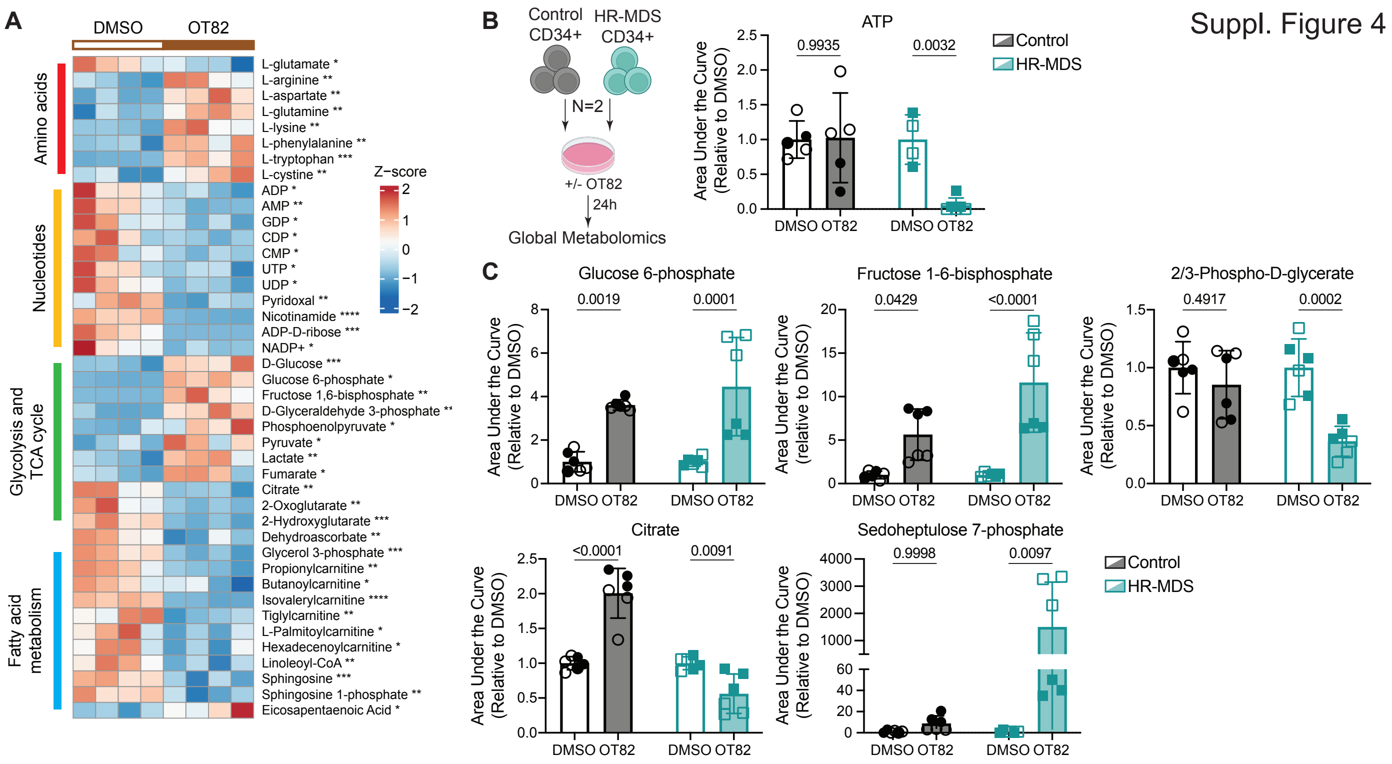


**Supplemental Figure 4: NAMPT inhibition disrupts metabolism in HR-MDS HSPCs**

**(A)** Heatmap comparing abundance of significantly expressed amino acids, nucleotides, glycolysis and TCA cycle metabolites and fatty acids in MDS-L cells treated with DMSO or 100nM OT-82 for 24h; measured using UHPLC/MS.

**(B)** Abundance of ATP measured using UHPLC/MS in HR-MDS (N=2) and healthy control (N=2) CD34+ HSPCs after 24h of treatment with DMSO or 100nM OT-82. Specimens used – M6-M7 and C6-C7.

**(C)** Abundance of Glucose-6-phosphate, Fructose-1,6-biphosphate, 2/3-phosphoglycerate, Citrate and Sedoheptulose-7-phosphate measured using UHPLC/MS in HR-MDS (N=2) and healthy control (N=2) CD34+ HSPCs after 24h of treatment with DMSO or 100nM OT-82. Specimens used – M6-M7 and C6-C7.

Suppl. Figure 5

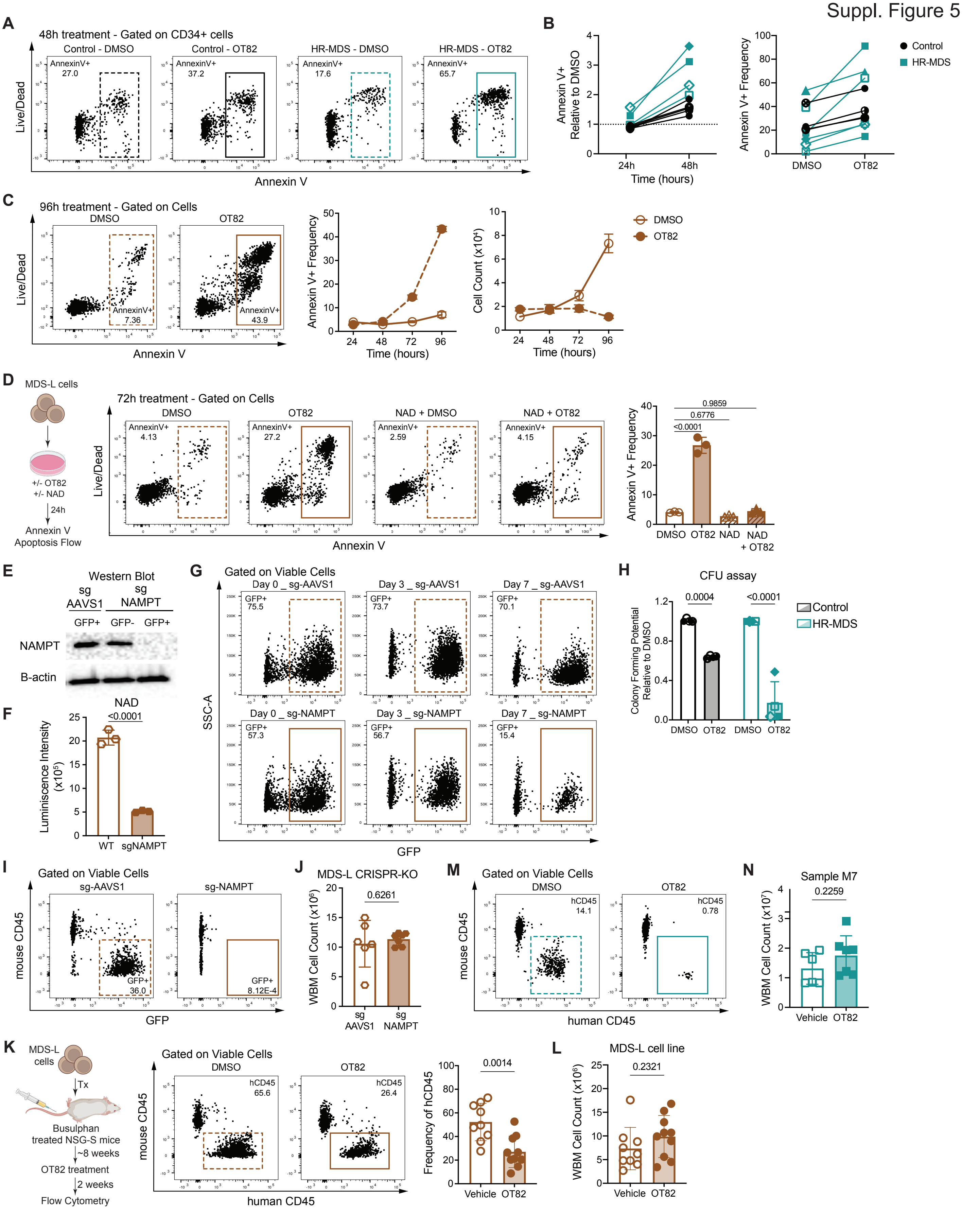

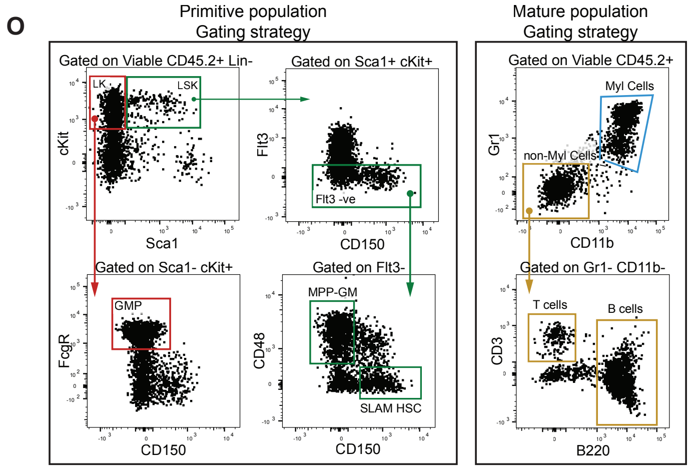


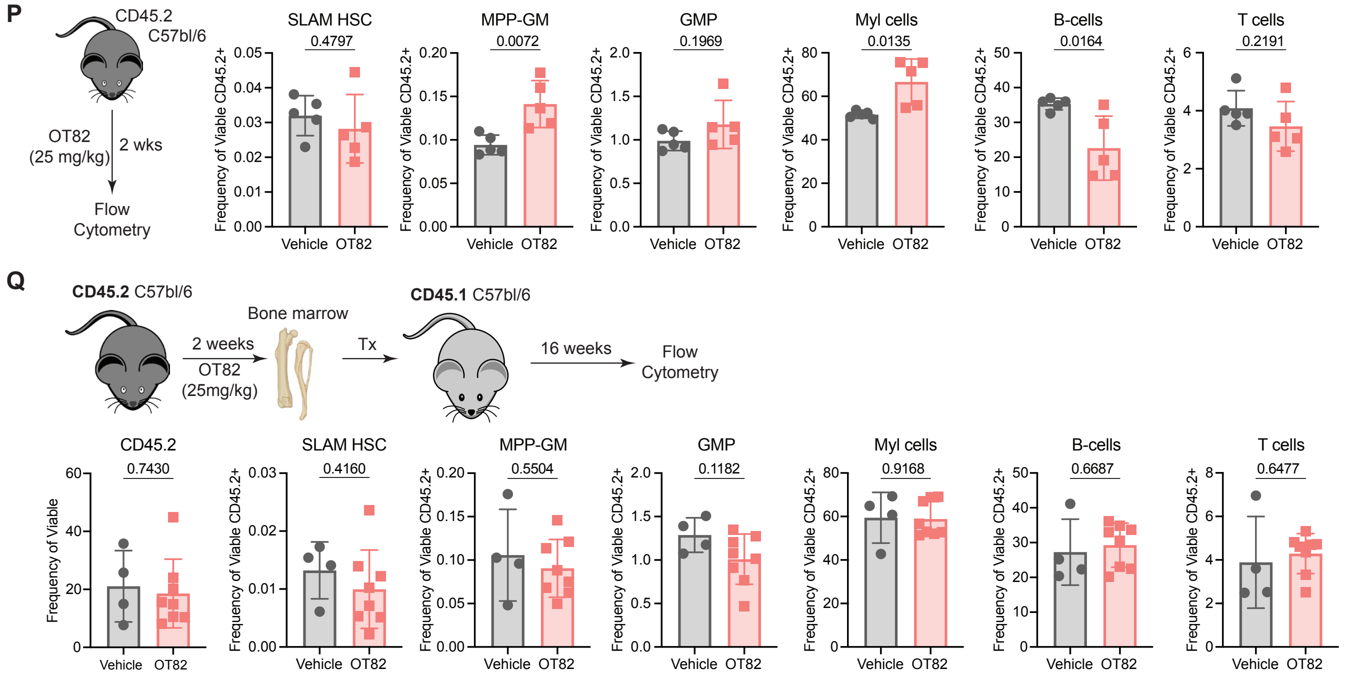


**Supplemental Figure 5: HR-MDS HSPCs are reliant on NAMPT for their survival and function as opposed to healthy control HSPCs**

**(A)** Representative flow plots showing Annexin V and Live/Dead gating strategy for HR-MDS (N=5) and healthy control (N=5) CD34+ HSPCs post 24h of treatment with DMSO or 100nM OT-82.

**(B)** Frequency of Annexin V+ cells relative to DMSO (left) and raw values (right) measured using flow cytometry in HR-MDS (N=5) and healthy control (N=5) CD34+ HSPCs post 48h of treatment with DMSO or 100nM OT-82. Specimens used – M6-M7, M9-12 and C3, C6-C9.

**(C)** Representative flow plots showing Annexin V and Live/Dead gating strategy (left), frequency of Annexin V+ cells bar pot (middle) and number of viable cells bar plot (right) for MDS-L cells post 24h of treatment with DMSO or 100nM OT-82.

**(D)** Representative flow plots showing Annexin V and Live/Dead gating strategy (left) and frequency of Annexin V+ cells bar plot for MDS-L cells post 24h of treatment with DMSO or 100nM OT-82 and supplemented with or without NAD.

**(E)** Western blot showing NAMPT expression post 24h transfection of MDS-L cells with sgRNA against NAMPT and AAVS1 control.

**(F)** NAD level measured using luminescence in MDS-L cells post 24h knock out with NAMPT and AAVS1 sgRNA.

**(G)** Representative flow plots showing GFP+ gating strategy for AAVS1 and NAMPT knockout MDS-L cells measured over time at day 0, day 3 and day 7.

**(H)** Number of colonies counted at day 12 for HR-MDS (N=4) and healthy (N=4) CD34+ HSPCs plated into CFU media post 24h of treatment with DMSO or 100nM OT-82. Each point is a biological replicate. Specimens used – M5-M8 and C7, C9-C11.

**(I)** Representative flow plots showing GFP+ gating strategy for hCD45 MDS-L cells in BM of NSG-S mice xenografts transplanted with AAVS1 and NAMPT knockout MDS-L cells.

**(J)** Whole bone marrow counts of NSG-S mice xenografted with sgAAVS1 and sgNAMPT MDS-L cells.

**(K)** Representative flow plots showing hCD45 gating strategy and frequency of hCD45+ cells in BM of NSG-S mice xenografted with MDS-L cells and treated with 25mg/kg OT-82 OG for 2 weeks (5 days on, 2 days off).

**(L)** Whole bone marrow counts of NSG-S mice xenografted with MDS-L cells and treated with 25mg/kg OT-82 OG for 2 weeks (5 days on, 2 days off).

**(M)** Representative flow plots showing hCD45 gating strategy for BM of NSG-S mice xenografted primary HR-MDS patient BM sample and treated with 25mg/kg OT-82 OG for 2 weeks. Specimen used – M7.

**(N)** Whole bone marrow counts of NSG-S mice xenografted with primary HR-MDS patient BM sample and treated with 25mg/kg OT-82 OG for 2 weeks (5 days on, 2 days off). Specimen used – M7.

**(O)** Flow cytometry plots depicting the gating strategy used for phenotypic identification of normal HSPC, myeloid and lymphoid populations in C57/B6 mice.

**(P)** Frequency of hematopoietic mature and immature populations in BM of C57/B6 mice treated with 25mg/kg OT-82 OG for 2 weeks (5 days on, 2 days off).

**(Q)** Frequency of hematopoietic mature and immature populations in BM of Jaxboy mice transplanted with BM collected from C57/B6 mice treated with 25mg/kg OT-82 OG for 2 weeks (5 days on, 2 days off).
